# Dual ecological-cultural pathways of sociocultural environment and resting state functional connectivity among Hispanic adolescents in the ABCD Study

**DOI:** 10.64898/2026.09.25.754508

**Authors:** Angela R. Laird, Chloe L. Hampson, Micaela Lembo, Katherine L. Bottenhorn, Kirthana Sukumaran, Carlos Cardenas-Iniguez, Erin Thompson, Shana Adise, Mariana Sanchez, Raul Gonzalez, Megan M. Herting, Alan Meca

## Abstract

Hispanic Americans face persistent health disparities rooted in complex sociocultural and bioecodevelopmental dynamics. Identifying early neurodevelopmental trajectories that confer risk or resilience during adolescence is critical for developing targeted health equity interventions. Grounded in the NIMHD Research Framework and bioecodevelopmental theory, data from 12- to 14-year-old Hispanic adolescents in the Adolescent Brain Cognitive Development (ABCD) Study were analyzed (n=236). Partial least squares correlation (PLSC) was used to evaluate multivariate associations between sociocultural measures—spanning individual, family, and community levels—and resting-state functional connectivity (rsFC) networks. *Post hoc* linear mixed-effects models examined how derived sociocultural-rsFC patterns predicted measures of physical and mental health, linking bioecodevelopmental patterns to adolescent well-being. PLSC revealed dual community- and family-level neural pathways of biological embedding, diverging from traditional nested bioecodevelopmental assumptions. The community pathway linked child opportunity and neighborhood disadvantage to connectivity across Cingulo-Opercular, Default, Dorsal Attention, Visual Networks. This community-rsFC profile was associated with lower respiratory functioning, reflecting an allostatic load marker of environmental stress. The family pathway linked caregiver ethnic identity and heritage acculturation to Default, Ventral Attention, Sensorimotor-Mouth, and Visual Networks connectivity, suggesting that the family system shapes neural systems underlying social attunement and identity formation. Structural community factors and familial culture impose distinct neurobiological demands on the adolescent connectome among Hispanic youth. Findings highlight respiratory health as a biological marker of environmental adversity and underscore the necessity of structural and culturally sustaining interventions to promote health equity among Hispanic youth.

## INTRODUCTION

Hispanic Americans face persistent health disparities across a spectrum of chronic conditions, including diabetes, hypertension, asthma, cancer, and obesity, as well as emotional and behavioral health (e.g., depression, suicide, substance use) ^1^. These disparities highlight an urgent need for a comprehensive understanding of Hispanic health that moves beyond adult symptomology to identify relevant neurodevelopmental trajectories that confer substantial risk for subsequent health concerns earlier in the lifespan. During adolescence, Hispanic (and non-Hispanic) youth experience dramatic changes in biopsychosocial development and engage in behaviors that exacerbate adverse health outcomes or promote positive health outcomes ^2^. Adolescence thus represents a critical ontogenic window where dramatic biopsychosocial shifts can either worsen health disparities or solidify trajectories toward health equity. Disentangling interrelations between adolescent development and risk/resilience factors necessitates a nuanced understanding of the social, cultural, and environmental determinants of health among Hispanic youth.

While prior work has examined individual sociocultural factors, such as cultural identity, values, and practices, a more integrated, holistic approach is required to capture the lived experience of Hispanic youth. We draw from bioecodevelopmental theory ^3^, as a conceptual framework for Hispanic adolescent brain function and development that emphasizes the multifaceted and multilevel nature of the sociocultural environment. Beyond traditional risk (e.g., economic disadvantage) and protective factors (e.g., family cohesion) across the broader ecological context, bioecodevelopmental theory posits that Hispanic youth also navigate a complex push/pull between cultural protectors (e.g., ethnic identity ^4^, familism ^5^) and cultural stressors (e.g., acculturation gap distress ^6^, perceived discrimination ^7^) ^3,8^. Critically, repeated or prolonged navigation of these chronic sociocultural stressors, unbuffered by protective factors, can trigger dysregulation across interconnected physiological systems (e.g., neuroendocrine, metabolic, immune, cardiovascular). This cumulative wear-and-tear on the body, termed allostatic load ^9^, serves as a key physiological mechanism linking chronic environmental stress to vulnerability for long-term physical and mental health conditions. Understanding how youth manage this stress response within their immediate environment is essential to identifying the early developmental trajectories of allostatic load.

In the current study, we empirically examined this conceptual framework using data from the Adolescent Brain Cognitive Development (ABCD) Study ^10^ and investigated the complex interplay between sociocultural environment, neurobiology, and risk/resilience factors for disparities among Hispanic youth. Grounded in the NIMHD Research Framework ^11^, we hypothesized that sociocultural variables at the individual, family, and community levels are critical contextual informants of the Hispanic adolescent environment that collectively predict neurobiological features associated with behavioral risk and resilience factors for health disparities. In this framework, central nervous system alterations represent front-line neural features of allostatic load. This work thus links knowledge across three domains of influence (i.e., sociocultural environment, (neuro)biological, and behavioral) and across three levels of influence (i.e., individual, interpersonal (family), and community) (Fig. 1). By modeling structural and intermediate social determinants ^12^ derived from responsible use of the ABCD Study dataset ^13^, we aimed to identify potential strategies for health equity interventions.

**Fig 1.**
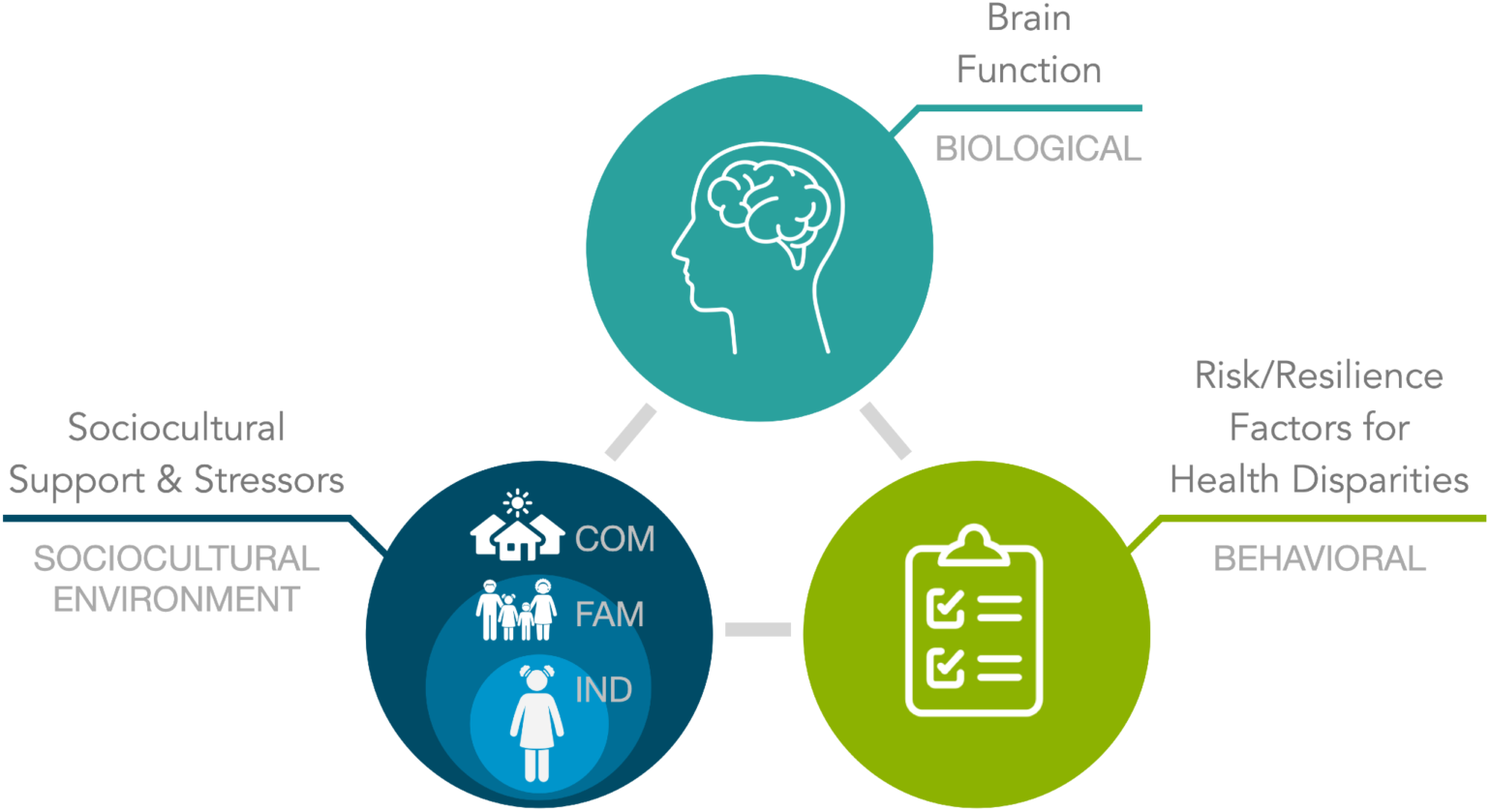
Conceptual Model. Adapted from bioecodevelopmental theory (Szapocznik and Coatsworth, 1999) and the NIMHD Research Framework. *Abbreviations*: Community (COM), Family (FAM), Individual (IND).

First, we applied partial least squares correlation (PLSC) ^14,15^ to identify latent, multivariate dimensions linking adolescent sociocultural environment (at the individual, family, and community levels) and resting state functional connectivity (rsFC). rsFC is an ideal neurobiological metric of interest given its sensitivity to cultural variables ^16–18^ and self-referential and affiliative processing ^19,20^. Because functional brain network organization mediates the central response to chronic environmental stress ^21–23^, rsFC variations offer a window into early neurobiological markers of allostatic processes. Second, for each significant latent dimension derived from PLSC, we determined how the identified sociocultural-brain pattern was related to indicators of physical (e.g., weight, physical inactivity, respiratory functioning, sleep) and mental health (e.g., internalizing/externalizing symptoms), thereby evaluating the developmental precursors that contribute to allostatic load and vulnerability to chronic health conditions later in the life course. The goals of this study were to examine bioecodevelopmental effects during adolescence and identify how individual, family, and community socioenvironment may differentially be associated with rsFC. By modeling structural and intermediate determinants, we aim to provide an evidence-based foundation for effective prevention and intervention strategies that extend beyond individual behavior change to address the roots of Hispanic health disparities.

## METHODS

### Participants

Participants were selected from the ABCD Study, the largest longitudinal study of brain development and child health in the United States (US) ^10^. Approximately 11,800 youth ages 9.00 to 10.99 years were enrolled in the ABCD Study across 21 sites in the US ^24^. Participants in the ABCD Study were recruited largely through geographically, demographically, and socioeconomically diverse school systems using epidemiologically informed methods to enroll a population-based, demographically diverse sample ^25^. Data from the ABCD Study were made available by the National Institute of Mental Health Data Archive (NDA). The current study used data from the ABCD Curated Annual Release 5.1 (http://doi.org/10.15154/z563-zd24) and primarily included data from the Year-3 and Year-4 Follow-Up when participants were 12-13 and 13-14 years old, respectively.

### Measures

#### Participant Demographics

Demographic information was provided at baseline by a child’s caregiver, including the child’s age, sex, ethnicity, and race, as well as the caregiver’s age, sex, ethnicity, race, education, and combined family income. In addition, caregivers reported the nativity (i.e., country of origin) for the child, parent/guardian, and grandparents ^26^.

#### Sociocultural Measures at the Individual Level

##### Youth Cultural Identity

Cultural identity is a sociocultural construct that refers to how individuals perceive, understand, and experience themselves in the context of a specific social group, racial background, or culture ^27^. Youth cultural identity was measured in the ABCD Study at the Year-3 Follow-Up using the youth-reported Multigroup Ethnic Identity Measure-Revised (MEIM-R) ^27,28^. The MEIM-R is a 6-item bidimensional measure with subscales that separately measure ethnic identity exploration (i.e., “*I have often talked to other people in order to learn more about my ethnic group*”) and commitment (i.e., “*I have a strong sense of belonging to my own ethnic group*”). The internal consistency is deemed ‘good’ according to Cronbach alpha values for exploration (*α* = 0.83) and commitment (*α* = 0.89).

##### Youth Cultural Practices

Acculturation refers broadly to the extent to which an individual retains their cultural heritage and/or adopts the receiving society’s culture ^29^ and has been hypothesized to play an important role in accounting for health disparities, particularly among Hispanic populations in the US ^30^. Youth acculturation practices were measured at the Year-3 Follow-Up using the youth-reported Vancouver Index of Acculturation (VIA). The VIA is a 16-item bidimensional measure with subscales that measure heritage acculturation (e.g., “*I often participate in my heritage cultural traditions*”) and US acculturation (e.g., “*I often participate in mainstream American cultural traditions*”) ^31^. Items address a range of cultural practices, including traditions, social activities, media, cultural values, and behavioral preferences. The internal consistency is high, with alpha values of 0.91 for heritage and 0.89 for US subscales.

##### Youth Cultural Values

Within Hispanic cultures, the family unit is often prioritized and significantly present, acting as the primary source of support, guidance, and identity ^32^. Youth cultural values were measured at the Year-4 Follow-Up using the Mexican American Cultural Values Scale (MACV) modified for youth ^32,33^. The ABCD-modified MACV is a 15-item measure with subscales that measure family support, family obligation, and family referent. Assessment of MACV youth-report data at the Year-2 Follow-Up indicated acceptable internal consistency for family support (*α* = 0.83), obligation (*α* = 0.77), and referent (*α* = 0.84) ^33^.

#### Sociocultural Measures at the Family Level

While prior work has demonstrated that acculturation and cultural identity are linked to maladaptive outcomes among Hispanic youth, including depression ^34,35^ and substance use ^36,37^, the underlying mechanisms of these associations are unclear. Some researchers have argued the need to conceptualize these effects within a family systems perspective ^38–40^, proposing that they are the result of sociocultural gaps (i.e., in differences in cultural identity, practices, or values) between Hispanic adolescents and their caregivers ^40–42^. These gaps may serve as significant stressors at the family level, leading to increased youth behavioral problems and substance use. For example, the acculturation discrepancy hypothesis posits that gaps in acculturation between caregivers and their adolescents may compromise family functioning, such as through poor communication or lack of involvement ^40^. Although sociocultural gaps are more common between immigrant caregivers and their US-born adolescents, studies have also documented sociocultural gaps persist among third or greater generation youth ^43^.

##### Caregiver Cultural Identity

Caregiver cultural identity was assessed using the caregiver-reported MEIM-R at the Year-3 Follow-Up. Good internal consistency was observed for caregiver ethnic identity exploration (*α* = 0.88) and commitment (*α* = 0.87) at the Year-2 Follow-Up ^33^.

##### Caregiver Cultural Practices

Caregiver cultural practices were assessed using the caregiver-reported VIA at the Year-3 Follow-Up. High internal consistency was observed for caregiver heritage acculturation (*α* = 0.92) and US acculturation (*α* = 0.91) subscales at the Year-2 Follow-Up ^33^.

##### Caregiver Cultural Values

Caregiver cultural values were assessed using the caregiver-reported MACV at the Year-4 Follow-Up. Acceptable internal consistency was observed for caregiver family support (*α* = 0.83), family obligation (*α* = 0.73), and family referent (*α* = 0.80) at the Year-2 Follow-Up ^33^.

#### Sociocultural Measures at the Community Level

The adolescent sociocultural environment is experienced both at home and beyond it. Neighborhood and broader community characteristics provide important contextualization of cultural, social, and physical ecological factors impacting child and adolescent development and play a role in the emergence of health inequities ^28,33,44,45^.

##### Perceived Discrimination

Perceived discrimination refers to self-reported perceptions of unfair treatment based on one’s ethnic or racial group ^46^ and prior research has indicated that 65% of Hispanic young adults ages 18-29 years have experienced or witnessed discrimination or unfair treatment because of their race or ethnicity ^47^. Perceived discrimination is negatively associated with a wide range of health outcomes ^48^; among children of color, exposure to racial/ethnic discrimination can be a chronic stressor and a risk factor for child health disparities ^49^. Youth perceived discrimination was measured in the ABCD Study at the Year-4 Follow-Up using the youth-reported Measure of Perceived Discrimination (MPD) ^33,50^. The MPD is a 7-item measure that includes questions about discrimination perceived to occur due to ethnic/racial background, including discrimination from teachers, adults outside of school, and other students, as well as feeling unwanted or unaccepted by “*American society*”. Acceptable internal consistency was observed for youth-reported perceived discrimination (*α* = 0.76) at the Year-2 Follow-Up ^33^.

##### Community Efficacy

Social cohesion and the capacity for collective self-regulation are key characteristics of neighborhoods ^51^ that are strongly associated with health-related problems, especially among adolescents ^52,53^. Neighborhood social cohesion was measured at the Year-4 Follow-Up using the caregiver-reported PhenX Neighborhood Collective Efficacy (CE) measure. The CE is a 10-item measure that includes two subscales of social cohesion and social control. Items related to social cohesion focus on trust in a community (e.g., “T*his is a close-knit neighborhood*”, “*People in this neighborhood can be trusted*”), while items related to social control attempt to capture neighbors’ willingness to intervene in the face of socially undesirable behaviors (e.g., “*If there was a fight in front of your house and someone was being beaten or threatened, how likely is it that your neighbors would break it up?*”) ^33^. Good internal consistency was observed for caregiver-reported social cohesion (*α* = 0.80) and social control (*α* = 0.83) at the Year-2 Follow-Up ^33^.

##### Linked External Data

The following three measures draw on geocoded information describing participants’ structural environment, derived from their primary residential addresses reported by their caregiver at the baseline timepoint when children were ages 9-10 years ^44,45^. The Child Opportunity Index (COI) 2.0 is a neighborhood composite measure that characterizes the community points of interventions relative to child health for the domains of education (e.g., early childhood centers, schools, educational and social resources), health and environment (e.g., toxic exposures, access to healthy food, green spaces), and social and economic (e.g., economic opportunities, economic and social resources) ^54^. The overall, nationally-normed COI child opportunity scores (i.e., from 1 to 100) were used ^44^. The NaNDA Neighborhood Disadvantage Index (NDI) is a measure of both neighborhood disadvantage, as well as neighborhood affluence and ethnic and immigrant concentration ^55^. The Getis-Ord Gi* Statistic measures residential segregation and emphasizes regions where racial/ethnic groups are concentrated or isolated^56^. The Gi* for Hispanic queen neighboring tracts was used ^44^.

#### Measures of Adolescent Health

Measures of adolescent physical and mental health were utilized as proximal markers of risk and resilience factors, serving as early-life predictors for the development of chronic disease in adulthood.

##### Anthropometrics and Blood Pressure

Height, weight, and blood pressure were assessed by a trained researcher; height and weight were measured annually ^26,57^, while blood pressure was assessed biannually starting at the Year-2 Follow-Up; for this study, we used data from the Year-4 Follow-Up. Youth were weighed to the nearest 0.1 inch and 0.1 pound up to three times, and the two closest measurements were averaged. Height and weight were converted into BMI (kg/m^2^). The CDC’s BMI percentiles were computed in SAS for descriptive clinical purposes only ^58^. Blood pressure was measured three consecutive times, with a 1-minute interval between each reading; the mean systolic and diastolic measurements were computed.

##### Physical Activity

Physical activity was assessed at the Year-4 Follow-Up using the youth-reported Youth Risk Behavior Survey ^59^. Participants responded to the item, “*During the past 7 days, on how many days were you physically active for a total of at least 60 minutes per day?*”.

##### Respiratory Functioning

Respiratory functioning was assessed at the Year-4 Follow-Up using a youth-reported measure of respiratory issues, including coughing/wheezing, allergies, and respiratory illness ^60,61^. Item-level questions included: (i) “*Have you ever had wheezing or whistling in the chest at any time in the past?*”, (ii) “*In the past 12 months, have you had a dry cough at night, apart from a cough associated with a cold or chest infection?*”, (iii) “*Has a doctor or other healthcare provider ever diagnosed you with asthma?*”, and (iv) “*Have you ever been diagnosed with pneumonia or bronchitis?*”.

##### Sleep

Hispanic adults report more sleep disturbances and consequences compared to their non-Hispanic peers; these sleep disparities are present among Hispanic children and adolescents ^62^ and have been linked to increased risk of chronic health conditions, including depression, substance use, and cardiovascular disease ^63^. Youth sleep was assessed using the youth-reported Munich Chronotype Questionnaire (MTCQ) ^64^ at the Year-4 Follow-Up. The MTCQ was used to compute a youth’s average weekly sleep duration and chronotype. Chronotypes describe an individual’s natural preference to wake and sleep at certain times; there are three chronotypes: morning, neither, and evening) ^65^, with evidence indicating that the evening chronotype is associated with psychiatric disorders ^66^.

##### Mental Health

Youth mental health was assessed at the Year-4 Follow-Up using the caregiver-reported Child Behavior Checklist (CBCL) ^67^, which assesses behavioral dimensions relevant to child psychopathology. Age- and gender-normed scores for internalizing and externalizing behaviors were used as measures of youth mental health ^57^.

### Hispanic Sample Selection

A total of 11,878 participants were enrolled in the baseline wave of the ABCD Study. In a prior study, we identified 2,411 baseline participants whose caregivers endorsed Hispanic/Latino/Latina ethnicity for the youth ^18^. For the present study, we evaluated longitudinal data from this Hispanic subgroup at the Year-3 and Year-4 Follow-Ups. From the original baseline cohort (N=2,411), sample size was reduced to a final analytic sample of N=236 due to a combination of longitudinal study retention across study waves and listwise deletion across incomplete participant datasets. Specifically, participants were retained only if they had youth- and caregiver-reported data at Year-2, Year-3, and Year-4 for the complete list of individual-, family-, and community-level sociocultural variables listed in Table 1. Descriptive statistics, including age, sex, and nativity, were calculated for the final analytic sample (N=236) and are presented in Table 2.

**Table 1.**
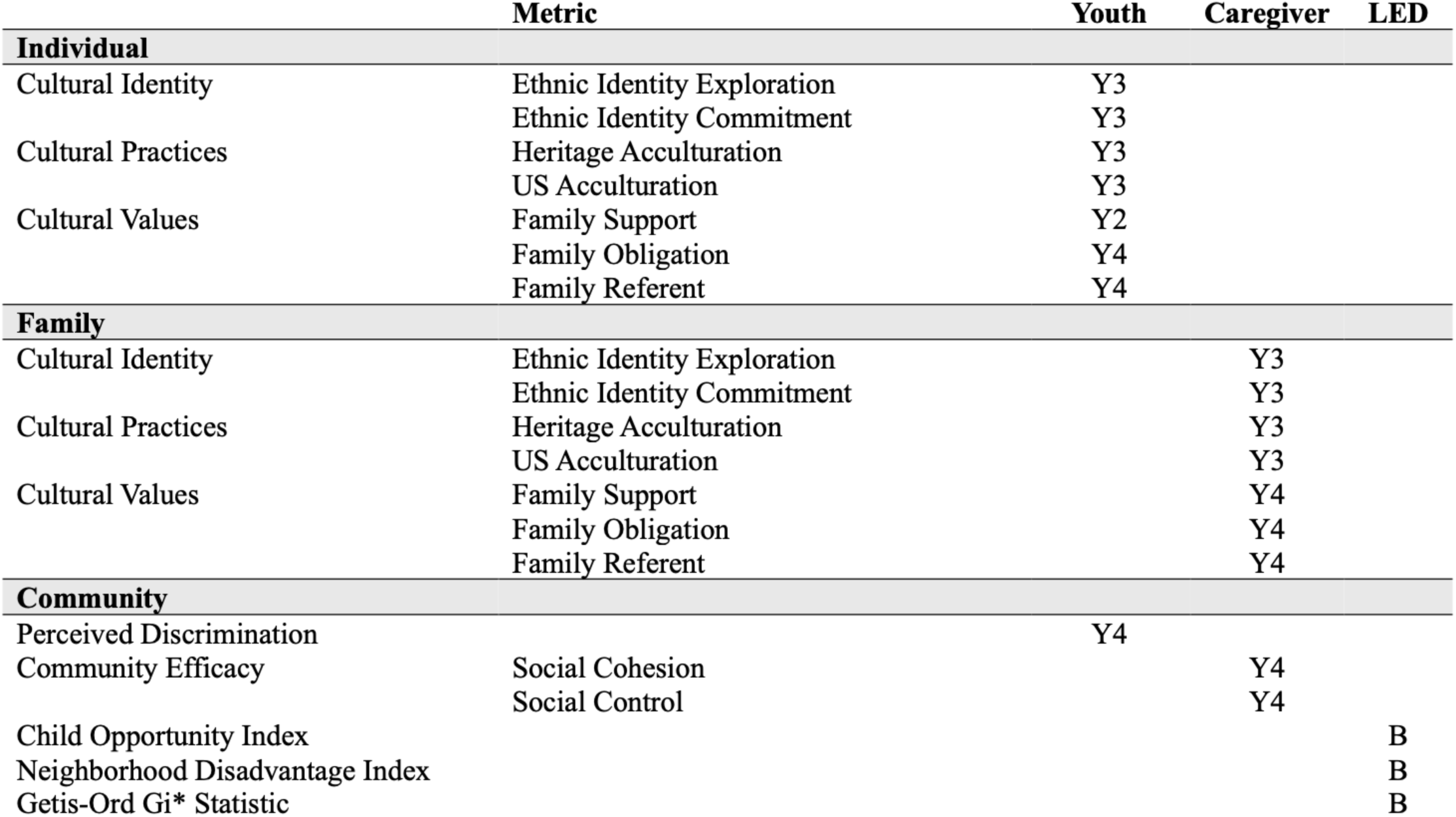
ABCD Measures of Sociocultural Environment. *Abbreviations*: Baseline (B), Year-2 Follow-Up (Y2), Year-3 Follow-Up (Y3), Year-4 Follow-Up (Y4).

|  | Metric | Youth | Caregiver | LED |
| --- | --- | --- | --- | --- |
| **Individual** |  |  |  |  |
| Cultural Identity | Ethnic Identity Exploration | Y3 |  |  |
|  | Ethnic Identity Commitment | Y3 |  |  |
| Cultural Practices | Heritage Acculturation | Y3 |  |  |
|  | US Acculturation | Y3 |  |  |
| Cultural Values | Family Support | Y2 |  |  |
|  | Family Obligation | Y4 |  |  |
|  | Family Referent | Y4 |  |  |
| **Family** |  |  |  |  |
| Cultural Identity | Ethnic Identity Exploration |  | Y3 |  |
|  | Ethnic Identity Commitment |  | Y3 |  |
| Cultural Practices | Heritage Acculturation |  | Y3 |  |
|  | US Acculturation |  | Y3 |  |
| Cultural Values | Family Support |  | Y4 |  |
|  | Family Obligation |  | Y4 |  |
|  | Family Referent |  | Y4 |  |
| **Community** |  |  |  |  |
| Perceived Discrimination |  | Y4 |  |  |
| Community Efficacy | Social Cohesion |  | Y4 |  |
|  | Social Control |  | Y4 |  |
| Child Opportunity Index |  |  |  | B |
| Neighborhood Disadvantage Index |  |  |  | B |
| Getis-Ord Gi* Statistic |  |  |  | B |

**Table 2.** Participants’ Demographic Characteristics. *N, n* = number of participants, *%* = percent of sample, *M* = mean, *SD* = standard deviation.

| **Youth (N = 236)** |  |
| --- | --- |
| **Youth Sex** | ***n* (%)** |
| Male | 134 (56.8%) |
| Female | 102 (43.2%) |
| **Youth Nativity** |  |
| US Born | 224 (94.9%) |
| Foreign Born | 12 (5.1%) |
| **Youth Age** | **Months / Years** |
| M (SD) Age | 168.04 (8.30) / 14.00 (0.69) |
| **Caregiver Sex** | ***n* (%) or *M* (SD)** |
| Male | 17 (7.2%) |
| Female | 219 (92.8%) |
| **Caregiver Age** | 38.84 yrs (5.81 yrs) |
| **Caregiver Education** |  |
| < High school diploma | 30 (12.7%) |
| High school diploma/GED | 37 (15.7%) |
| Some college | 34 (14.4%) |
| Associate's degree | 45 (19.1%) |
| Bachelor's degree | 49 (20.8%) |
| Postgraduate degree | 41 (17.4%) |
| **Household Income per Year (\$US)** | |
| <\$12,000 | 19 (8.1%) |
| \$12,000-\$15,999 | 10 (4.2%) |
| \$16,000-\$24,999 | 13 (5.5%) |
| \$25,000-\$34,999 | 22 (9.3%) |
| \$35,000-\$49,999 | 44 (18.6%) |
| \$50,000-\$74,999 | 42 (17.8%) |
| \$75,000-\$99,999 | 44 (18.6%) |
| \$100,000 and greater | 42 (17.8%) |

### Neuroimaging Data

Youth participants completed a neuroimaging protocol at the Year-4 Follow-Up that included structural MRI and rs-fMRI using high spatial and temporal resolution simultaneous multislice/multiband echo-planar imaging (EPI) ^68,69^. For Siemens scanners, fMRI scan parameters were as follows: 90 x 90 matrix, 60 slices, field of view = 216 x 216, echo time/repetition time = 30/800 ms, flip angle = 52°, 2.4-mm isotropic resolution, and slice acceleration factor 6. The complete protocols for all vendors and sequences have been provided by Casey et al. ^68^. Participants were scanned while they completed four 5-minute resting-state BOLD fMRI scans with their eyes open and fixated on a crosshair.

Imaging data processing was performed by the ABCD Data Analysis, Informatics, and Resource Center (DAIRC) ^69^. Measures of rsFC are based on a seed-based correlational approach adapted for cortical surface-based analysis. Processed rsFC data include the tabulated results of region of interest (ROI)-based analyses. Following pre-analysis processing steps (i.e., removal of initial frames, normalization, regression, temporal filtering, and calculation of ROI-average time course), imaging data were preprocessed through surface sampling and ROI averaging. Then, correlation values were calculated for each pair of ROIs, which were Fisher transformed to normally distributed z-values and averaged within or between networks to provide summary measures of network correlation strength. Fisher Z-transformed averages of all pairwise correlations within each of the Gordon networks ^70^ were made available. The Gordon networks included in this analysis were: Cingulo-Opercular (CON), Cingulo-Parietal (CPN), Default (DN), Dorsal Attention (DAN), Fronto-Parietal (FPN), Retrosplenial-Temporal (RTN), Salience (SN), Sensorimotor-Hand (SHN), Sensorimotor-Mouth (SMN), Ventral Attention (VAN), and Visual (VN). We examined both within- (e.g., DN-DN) and between-network (e.g., CON-DN, DN-FPN) rsFC correlations.

### Analyses

The goals of the current study were to first examine how sociocultural environment at the individual, family, and community levels was associated with rsFC among Hispanic ABCD youth, and then evaluate how the identified sociocultural-rsFC patterns were associated with physical and mental health measures that serve as risk/resilience factors for subsequent chronic health conditions. Figure 2 provides an overview of the analytic approach.

**Fig. 2.**
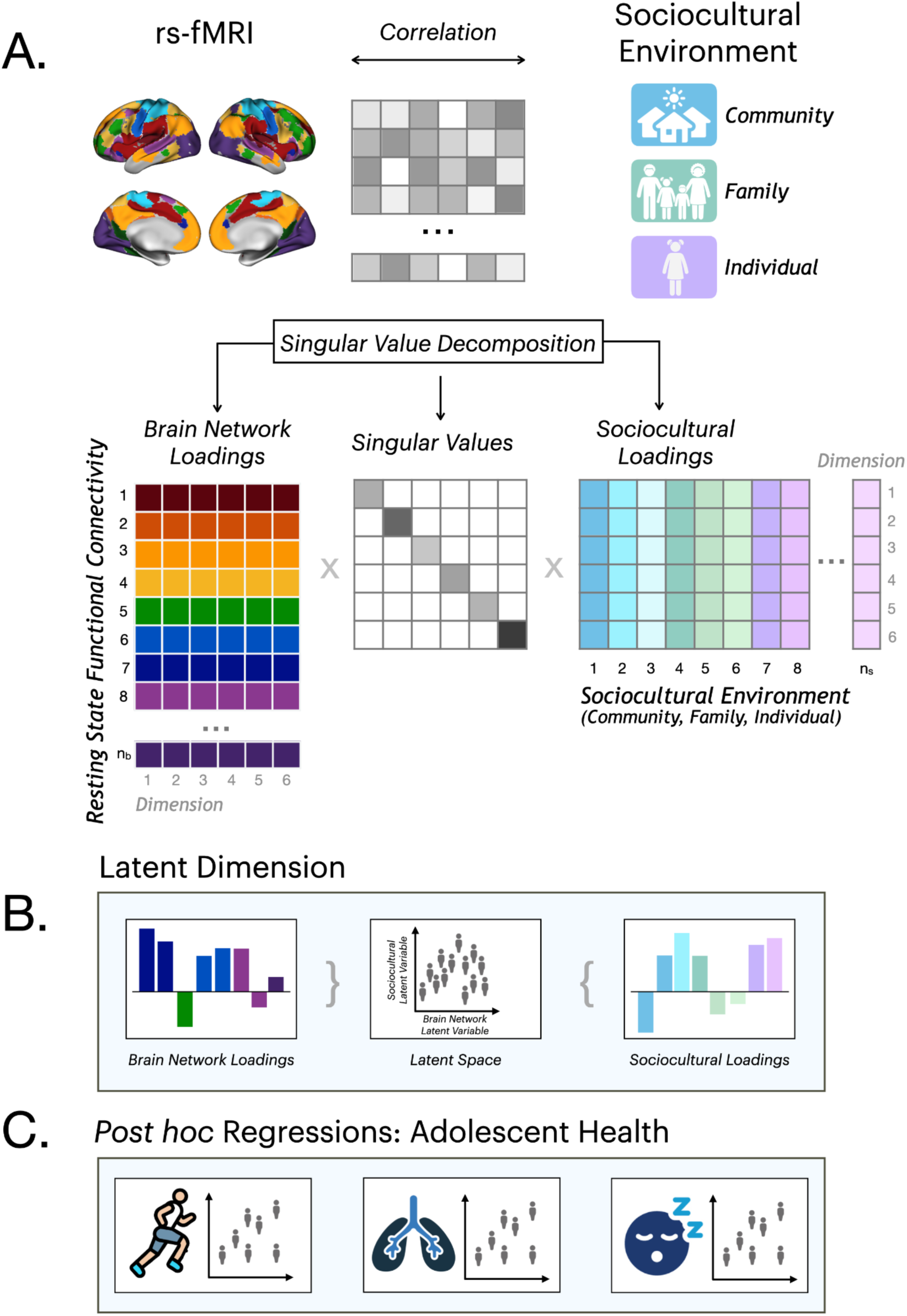
Analytic Approach Overview. (A) Partial least squares correlation (PLSC) identified shared information between resting state functional magnetic resonance imaging (rs-fMRI) data and sociocultural environment variables. Singular value decomposition of the correlation matrix for the two data blocks yielded three matrices, including a latent matrix for brain network loadings, the singular values, and a latent matrix for sociocultural loadings. (B) A latent dimension and its associated two latent variables (one per data block). (C) *Post hoc* regressions were performed to determine associations between measures of adolescent health (e.g., physical activity, respiratory functioning, sleep) and participant-level contributions to the latent dimensions identified by the PLSC analysis.

### Partial Least Squares Correlation

Descriptive statistics were computed across the final sample, including mean and standard deviation for all continuous variables. We applied partial least squares correlation (PLSC), a multivariate statistical approach that analyzes the relations between two multidimensional blocks of variables (here, sociocultural environment and rsFC) and reveals shared covariance patterns ^14,15^. PLSC identifies latent dimensions that best capture associations between the variable blocks while maximizing their covariance. PLSC requires complete data; thus, participants with missing data were removed using listwise deletion. PLSC was performed on the variables for sociocultural environment (Table 1) and Gordon networks.

Following the work of Sukumaran et al. ^71^ and Bottenhorn et al. ^72^, sociocultural and rsFC variables were residualized, mean-centered, and normalized to a standard deviation of one to remove the influence of demographic, scanner-related, and confounding influences (i.e., youth age and sex; caregiver age, sex, and education; household income; site; scanner manufacturer; head motion). Singular value decomposition (SVD) of the cross-correlation matrix between the two sets/blocks of variables.yielded sets of matrices describing: (i) variable loadings for rsFC measures, (ii) variable loadings for sociocultural measures, and (iii) all the singular values. Each latent dimension consists of two paired latent variables derived from linear combinations of the original sociocultural and rsFC data. The percentage of variance explained by each latent dimension relative to other latent dimensions was computed. Permutation testing using 10,000 permutations was performed to determine the statistical significance of the global model and each latent dimension. To determine the significance of the specific variable loadings, we computed standard error estimates using a bootstrap procedure (n = 10,000 samples), which yielded a confidence bootstrap ratio. As bootstrap ratios are analogous to Z-scores, loadings with bootstrap ratios greater than 2.5 (corresponding to P < 0.01) were considered reliable and significant. As PLSC involves a single decomposition (i.e., one test per latent dimension), multiple comparisons corrections were not implemented.

PLSC analysis was conducted in `R` (version 4.5.2). We used the `TExposition` package ^73^, followed by permutation testing using the `perm4PLSC()` function to determine the number of significant latent dimensions; the bootstrap validation, which assesses the stability of the variable contribution to each latent dimension, was executed with the `Boot4PLSC()` function in package `data4PCCAR` (https://github.com/HerveAbdi/data4PCCAR).

#### Associations with Adolescent Health

Next, we conducted *post hoc* regression analyses to determine associations between physical and mental health measures (Table 3) and participant-level contributions to the identified latent dimensions. We implemented linear mixed effect models adjusted for confounding and precision covariates listed above (i.e., youth age and sex; caregiver age, sex, and education; household income; site; scanner manufacturer; head motion), with random effects included to control for the structure of the sample including study site and family relationship.

**Table 3.** ABCD Measures of Adolescent Physical and Mental Health. *Abbreviations*: Year-4 Follow-Up (Y4).

|  | Metric | Youth | Caregiver |
| --- | --- | --- | --- |
| **Physical Health** |  |  |  |
| Height and Weight | BMI | Y4 |  |
| Blood Pressure (Systolic and Diastolic) | Mean Arterial Blood Pressure | Y4 |  |
| Physical Activity | Youth Risk Behavior Survey | Y4 |  |
| Respiratory Functioning | Wheezing/Whistling, Dry Cough,<br>Asthma, Pneumonia/Bronchitis | Y4 |  |
| Sleep | Average Weekly Sleep Duration | Y4 |  |
|  | Chronotype | Y4 |  |
| **Mental Health** |  |  |  |
| Child Behavior Check List | Internalizing Problems |  | Y4 |
|  | Externalizing Problems |  | Y4 |

## RESULTS

Three significant latent dimensions were identified in the multivariate associations between adolescent sociocultural environment and resting state functional connectivity (<u>Fig. S1</u>). First, a significant latent dimension that explained 24.27% of the variance common to these two sets of variables was observed. Based on the brain network loadings (i.e., rsFC correlations) and sociocultural loadings (i.e., sociocultural variables) that met a bootstrap ratio of 2.5 (*P* < 0.01), a complex pattern of association was observed for the neighborhood-level sociocultural measures of Child Opportunity Index (COI) and NaNDA Neighborhood Disadvantage Index (NDI) and CON, DN, DAN, and VN connectivity (Fig. 3). Specifically, COI (i.e., opportunity) was *positively* and NDI (i.e., disadvantage) was *negatively* associated with within-network connectivity for the DN and VN. Furthermore, COI was *negatively* and NDI was *positively* associated with DN connectivity with CON, DAN, and VN.

**Fig. 3.**
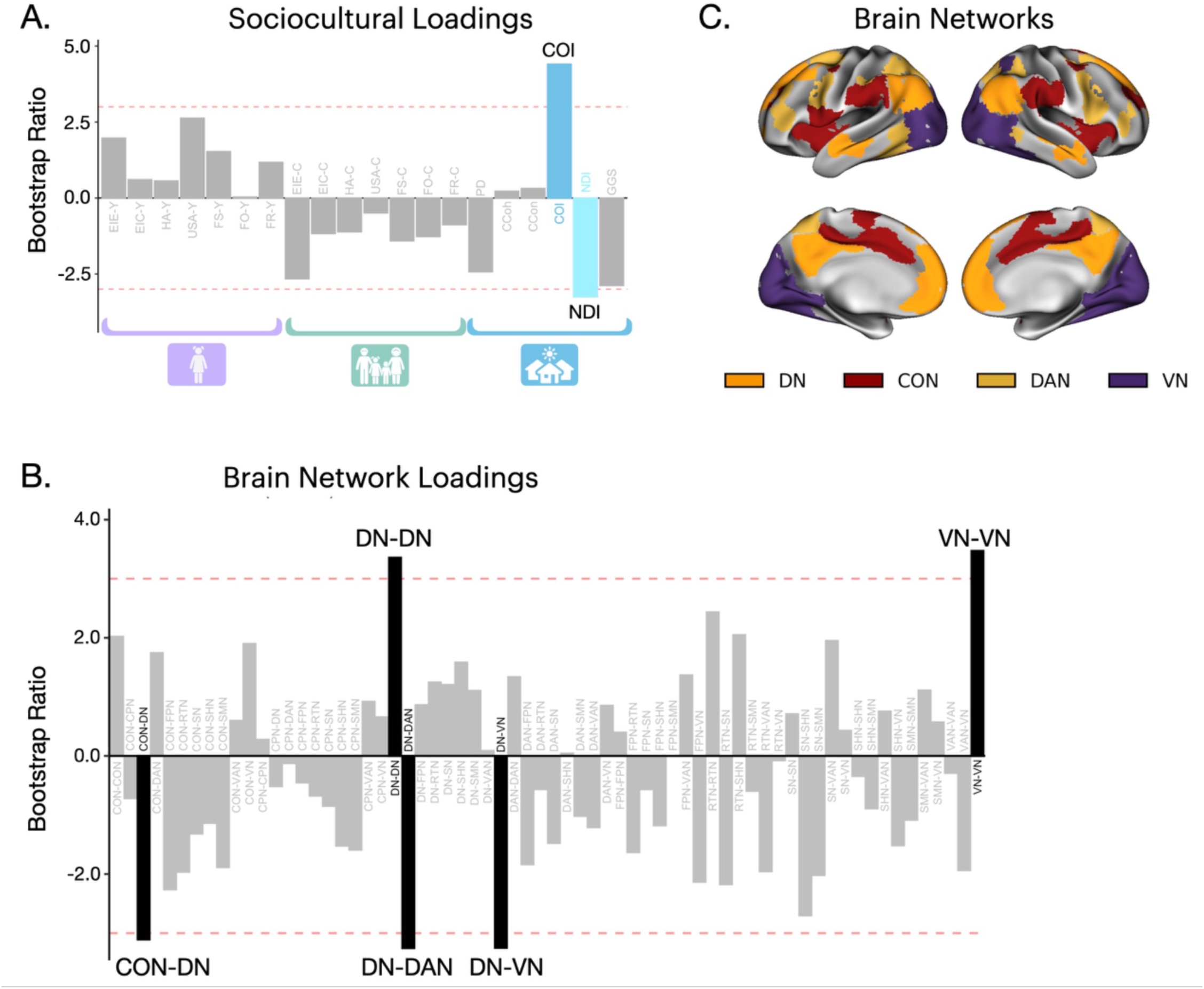
Latent Dimension 1 Results. Bootstrap ratios for (A) sociocultural variables and (B) rsFC brain network loadings for latent dimension (LD) 1. (C) Lateral and medial surfaces for LD1 brain networks. Blue (community-level sociocultural) or black (brain network) bars indicate a bootstrap ratio of 2.5 (*P* < 0.01). *Abbreviations*: Child Opportunity Index (COI), NaNDA Neighborhood Disadvantage Index (NDI), Cingulo-Opercular Network (CON), Default Network (DN), Dorsal Attention Network (DAN), Visual Network (VN).

For the second significant latent dimension (15.91% variance explained), bootstrap ratios revealed that caregiver-report family obligation (FO-C) and family referent (FR-C) from the MACV measure were the most influential variables (Fig. 4A). However, bootstrap ratios for the rsFC correlations were not stable (Fig. 4B). That is, we observed a significant negative association between caregiver cultural values and brain connectivity, but we were unable to determine which brain network(s) reliably contributed to this pattern.

**Fig. 4.**
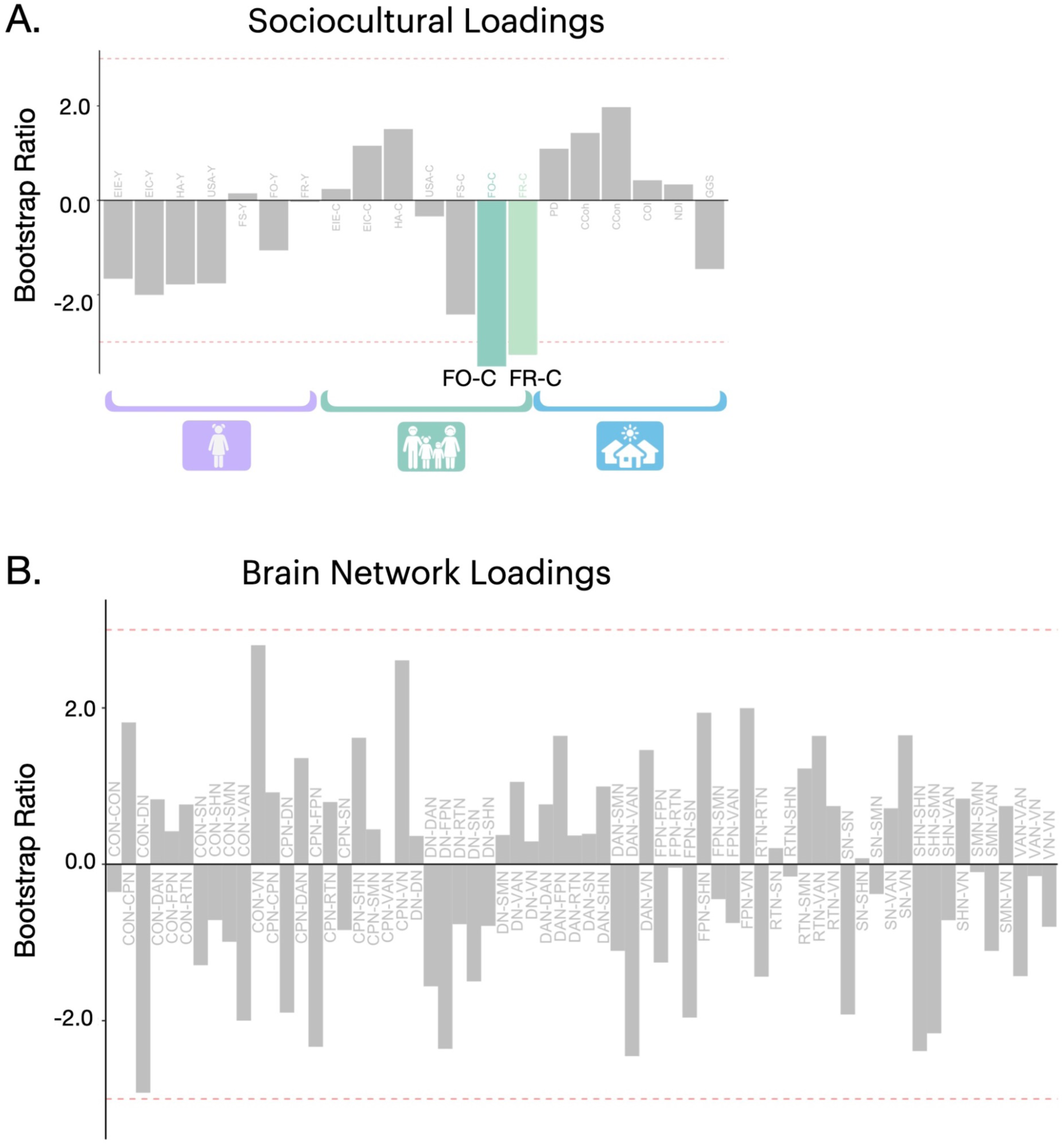
Latent Dimension 2 Results. Bootstrap ratios for (A) sociocultural variables and (B) rsFC brain network loadings for LD2. Green (family-level sociocultural) bars indicate a bootstrap ratio of 2.5 (*P* < 0.01). *Abbreviations*: caregiver-reported Family Obligation (FO-C) and Family Referent (FR-C).

For the third significant latent dimension (12.98% variance explained), bootstrap ratios showed that caregiver ethnic identity exploration (EIE-C) and commitment (EIC-C), as well as caregiver heritage acculturation (HA-C) were negatively associated with DN-SMN and VAN-VN between-network connectivity (Fig. 5).

**Fig. 5.**
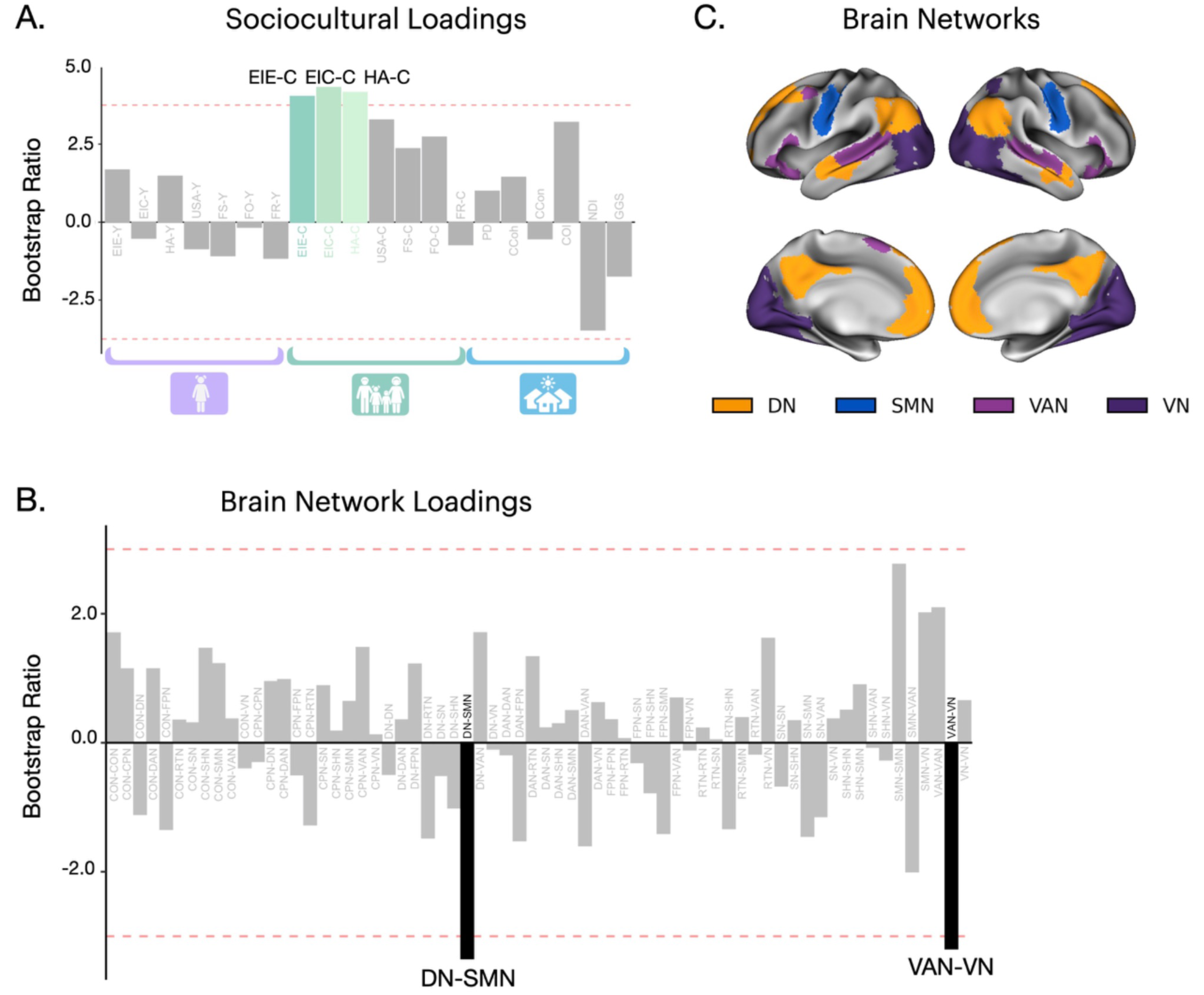
Latent Dimension 3 Results. Bootstrap ratios for (A) sociocultural variables and (B) rsFC brain network loadings for LD3. (C) Lateral and medial surfaces for LD3 brain networks. Green (family-level sociocultural) or black (brain network) bars indicate a bootstrap ratio of 2.5 (*P* < 0.01). *Abbreviations*: caregiver-reported Ethnic Identity Exploration (EIE-C), caregiver-reported Ethnic Identity Commitment (EIC-C), caregiver-reported Heritage Acculturation (HA-C), Default Network (DN), Sensorimotor-Mouth Network (SMN), Ventral Attention Network (VAN), Visual Network (VN).

Consistently across latent dimensions LD1 and LD3, protective factors (i.e., opportunity, caregiver identity) were negatively associated with between-network connectivity, while risk factors (i.e., disadvantage) were positively associated with within-network connectivity.

Linear mixed effects modeling revealed significant associations between measures of adolescent health and individual differences in identified latent dimensions, as well as rsFC correlations. For the first latent dimension, the multivariate pattern for neighborhood-level sociocultural measures (i.e., COI and NDI) and CON, DN, DAN, and VN connectivity was associated with respiratory issues, specifically wheezing/whistling (P = 0.037) (Fig. 6); however, this association did not survive correction for multiple comparisons. The second latent dimension was not analyzed given the unstable rsFC bootstrap ratios. No significant health measures were observed for the third latent dimension. The complete set of results for the first and third latent dimensions is shown in <u>Table S1</u>.

**Fig. 6.**
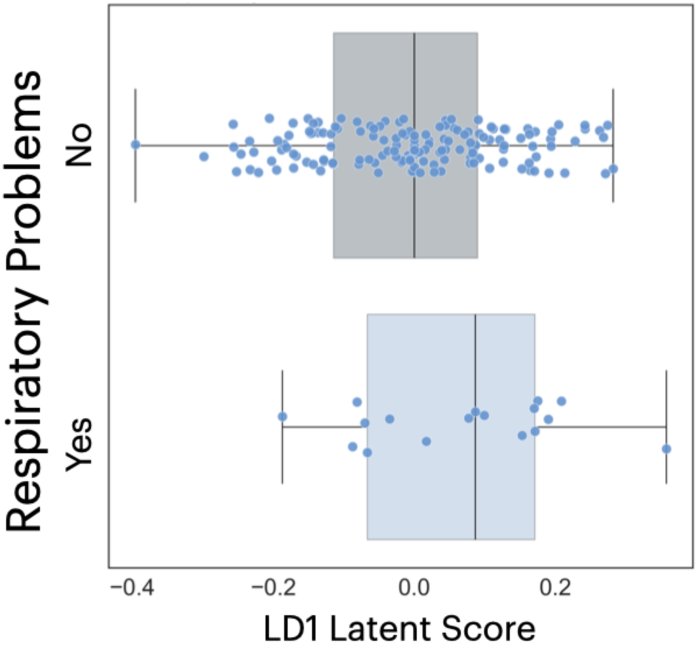
Associations with Adolescent Health. For LD1, the multivariate pattern for neighborhood (COI and NDI) and connectivity (CON, DN, DAN, and VN) was associated with elevated respiratory issues (i.e., *wheezing/whistling*) (*P* = 0.037). *Abbreviations*: Child Opportunity Index (COI), NaNDA Neighborhood Disadvantage Index (NDI), Cingulo-Opercular Network (CON), Default Network (DN), Dorsal Attention Network (DAN), and Visual Network (VN).

## DISCUSSION

A robust, multivariate statistical technique was applied to identify latent associations between youth sociocultural environment and resting state functional connectivity among over 200 Hispanic adolescents ages 12-14 years enrolled in the ABCD Study. Macro-ecological determinants, specifically child opportunity and neighborhood disadvantage, were associated with key large-scale neurocognitive networks. These results suggest that the community context serves as a distal scaffold for childhood development of the Cingulo-Opercular (CON), Default (DN), Dorsal Attention (DAN), and Visual (VN) Networks, potentially mediating relations between environmental quality and complex human cognition. Next, caregiver cultural identity and heritage acculturation were associated with connectivity across a diverse set of neural systems, including the DN, Sensorimotor-Mouth Network (SMN), Ventral Attention Network (VAN), and VN. These correlations may reflect how the family microsystem acts as a critical medium for biological embedding, where transmission of cultural identity and practices influences the neural architecture underlying self-referential thought, social salience, and sensory-motor integration.

### Theoretical Divergence: Decoupling Community and Family Systems

Bioecodevelopmental theory ^3^ is a framework designed to understand and prevent problem behaviors (e.g., substance use problems, delinquency) by examining a child through the lens of their entire social ecosystem. Under the bioecodevelopmental paradigm, behavioral outcomes are viewed as the product of transactional influences across nested ecological strata. Within these systems, the interplay between distal and proximal risk factors and buffering protective mechanisms determines the developmental trajectory of the individual. Importantly, as argued by Szapocznik and Coatsworth ^3^ and others ^74,75^, family is the most proximal environment to the child (i.e., the site of the most frequent and intense reciprocal interaction) as well as the primary vehicle for cultural socialization ^76^; therefore, family is the central mechanism of development for the child’s biological and psychological self-regulation. In contrast, the current results suggest that community plays a stronger, more prominent role, operationalized through the provision of resources, norms, social safety nets, as well as the prevalence of stressors. Whereas bioecodevelopmental theory views community and family as interconnected, with community as the distal ecological context that acts as a moderator of family processes, we observed separate latent dimensions for community- and family-level variables. That is, rather than nested, interacting strata, the current results suggest community and family are associated with different neurobiological systems of development. This suggests a divergent embedding process, where structural community factors and familial culture impose distinct, non-redundant demands on the developing connectome.

### A Community-Level Ecological Pathway

Results from the first latent dimension indicated that child opportunity and neighborhood disadvantage represent critical community-level contextual stressors that influence neurodevelopmental trajectories. The observed associations across the “Triple Network” (i.e., CON, DN, DAN) ^77^ and the Visual Network suggest that these neighborhood-level factors may calibrate the brain’s ability to toggle between internally-directed cognition and externally-directed sensory processing. This reflects a neurobiological adaptation to the safety and resource density of the child’s immediate ecosystem; such a mechanism likely operates across human development, though its specific phenotypic expression may interact with culturally patterned environmental assets and stressors. In our sample, the association between community contexts and altered DAN and DN connectivity suggests environmental calibration of attentional control systems. For instance, chronic exposure to neighborhood-level disadvantage may necessitate a shift toward stimulus-driven, hyper-vigilant processing of the CON ^78–81^ at the expense of top-down, goal-directed regulation (i.e., attentional control via the DAN ^82–84^ and internal reflection via the DN ^20^. Consequently, while these findings illustrate how the community environment functions as a biological embedding mechanism translating socioeconomic strata translate into divergent patterns of functional system, future research is needed to determine whether these specific neural profiles generalize across racially and ethnically diverse cohorts or reflect unique socio-ecological dynamics among Hispanic youth.

### A Family-Level Cultural Pathway

The third latent dimension suggested that familial cultural ontogeny significantly shapes adolescent brain architecture. Observed associations between caregiver identity/acculturation and connectivity in the VAN, which is responsible for stimulus-driven attention, and the DN, which is involved in internal mentation, suggests that family cultural schemas may modulate how adolescents prioritize environmental cues relevant to the development of their self-identity. Indeed, prior work has found important associations between caregiver and adolescent cultural identity among recently immigrated middle-to-late Hispanic adolescents ^85^. The interconnection between these parents and adolescents may stem from internalization of overt and/or covert ethnic socialization ^76^. Furthermore, the involvement of the SMN and VN may reflect neurobiological signatures of culturally-specific communication patterns and social-perceptual styles. That is, regions associated with speech and motor coordination and visual processing may influence systems involved in interpersonal expression and social observation. Collectively, these results provide evidence that the family microsystem provides a culturally-coded environment that exerts a top-down influence on both associative and primary cortical connectivity. While these observational associations require prospective validation, they highlight the potential biological relevance of heritage culture retention within the family microsystem. Notably, these neurobiological patterns align with decades of historical, anthropological, and developmental research demonstrating that cultural retention and ethnic socialization serve as vital protective scaffolds during youth development ^74,86,87^. In a broader socioecological context, if heritage culture retention supports functional network integration, policies or social pressures that marginalize heritage languages and traditions (i.e., “*assimilationist*” policies) could inadvertently disrupt key protective developmental environments. Conversely, supporting environments that foster biculturalism and bicultural competence, which are consistently linked to positive psychological adjustment ^88,89^, may align with and support adaptive neural plasticity as adolescents navigate complex multicultural environments.

### Respiratory Functioning as an Indicator of Systemic Adaptation to Environmental Determinants

The current study also identified a multivariate association between community-level determinants, large-scale neural networks, and adolescent respiratory functioning. Consistent with the framework of allostatic load ^90,91^, these data indicate that the respiratory system may act as a physiological marker for the neural impact of community-level stressors. These results suggest that child opportunity and neighborhood disadvantage serve as distal drivers of a neurobiobehavioral profile characterized by the co-regulation of the CON, DN, DAN, and VN alongside pulmonary efficiency and suggests a systemic adaptation to environmental pressure. Lower respiratory functioning—often a proxy for chronic stress or environmental pollutants ^92^—covaries with neural shifts toward stimulus-driven processing and altered executive control. This highlights a bottom-up mechanism where the physiological toll of a disadvantaged community environment is reflected in the functional integration of the brain’s primary cognitive control centers.

### Implications for Culturally-Competent Interventions

Mapping of community and family culture onto distinct neural architectures provides a robust empirical foundation for the development of multisystemic, culturally-attuned interventions. Our findings suggest that among Hispanic adolescents, the functional connectivity of CON, DN, and DAN networks is significantly calibrated by the macro-ecological context of child opportunity and neighborhood disadvantage. This suggests that clinical efforts to improve executive function must look beyond individual cognitive training and address the structural disadvantage that necessitates high-arousal, stimulus-driven neural states. Importantly, we emphasize that not all sociocultural factors are malleable, particularly those at the community level. For example, residential segregation should not be framed as “*housing decisions that people make*” but rather “*housing decisions constrained by structural factors*”. Furthermore, the association between caregiver cultural identity and heritage acculturation and VAN and SMN networks highlights the family microsystem as a critical lever for neurobiological regulation. Effective interventions should prioritize restructuring family interactions to reinforce protective cultural schemas that buffer against neighborhood-level risk. By aligning clinical goals with an adolescent’s neuro-cultural signature, practitioners can foster a systemic fit that leverages familial strengths, such as familismo ^93^, to recalibrate the attentional and sensory systems taxed by environmental adversity. Ultimately, these results suggest that among Hispanic youth, promoting an individual’s well-being requires a dual-focus on expanding community-level opportunity while fortifying the cultural integrity of the family unit.

### Limitations

While the current findings provide compelling evidence for the biological embedding of sociocultural factors, several limitations regarding data scope, measurement, and generalizability must be considered. First, although the ABCD Study includes a large, demographically diverse cohort ^25^, our analytic sample was restricted to a smaller subset of participants whose caregivers endorsed Hispanic youth ethnicity ^18^. This reduction in sample size was primarily driven by listwise deletion across the multi-domain set of individual-, family-, and community-level sociocultural variables, which were collected over three waves of the ABCD Study and included both youth- and caregiver-reports. Thus, participant inclusion in our sample required both longitudinal study retention across multiple waves and within-wave protocol completion of the relevant measures. Reliance on caregiver reports to define youth ethnicity represents a key measurement limitation, as early-to-middle adolescents are actively negotiating their own ethnic-racial identity; future work should incorporate youth self-identification as this information becomes available across later ABCD waves (e.g., Year-8 Follow-Up). Furthermore, our sample reflects a specific sociocultural context that may not generalize across all adolescents, or even across distinct Hispanic subgroups (e.g., Mexican American vs. Cuban American) and global populations (e.g., Central vs. South American) who encounter unique historical, sociopolitical, and economic pressures. Relatedly, we were unable to fully account for youth or family immigration status and documentation history, which serve as critical structural factors shaping acculturative stress, legal vulnerability, and healthcare access.

Second, our observational, cross-sectional design restricts our capacity to draw causal inferences or determine the precise directional dynamics between environmental exposures, cultural processes, and functional connectivity patterns. Given that data were restricted to a narrow ontogenetic window (ages 12–14 years) and some community-level exposures were measured at baseline (ages 9–10 years), the observed multivariate associations likely reflect a transient phase of maturational sensitivity rather than static neural architecture. Indeed, default mode ^94,95^ and attentional ^96,97^ networks undergo extensive synaptic pruning and myelination throughout later adolescence when ethnic identity processes become particularly developmentally salient ^88,98^. Future longitudinal work leveraging subsequent ABCD waves will be essential to assess time-varying exposures and test directional, lagged developmental pathways. Finally, although the ABCD Study includes a wealth of variables across ecological contexts, measurement of key cultural stressors relevant for Hispanic youth were not included (e.g., bicultural stress, intragroup marginalization ^99^), limiting our capacity to fully model the dynamic interplay between cultural risk and protective factors.

## CONCLUSIONS

The present study provides an integrated mapping of how macro-level ecological determinants, familial cultural ontogeny, and somatic physiology converge to shape the functional architecture of the Hispanic adolescent brain. By demonstrating that community-level variables (i.e., child opportunity and neighborhood disadvantage) and family-level variables (i.e., caregiver ethnic identity and heritage culture retention) correlate with distinct yet overlapping neural networks, these findings argue for a multisystemic view of neurodevelopment. Specifically, the results suggest a dual-pathway for biological embedding. First, our findings provide evidence of an ecological pathway in which community-level opportunity and neighborhood disadvantage act as distal scaffolds that calibrate the CON, DN, DAN, and VN. This pathway is uniquely coupled with respiratory functioning, suggesting that environmental disadvantage imposes a systemic allostatic load ^100^ that synchronizes pulmonary and neural regulation to navigate high-stress ecosystems. Second, we demonstrated a cultural pathway in which family-level ethnic identity and heritage culture retention serve as a proximal filter and regulate the SMN and VAN. This indicates that family culture provides a specialized sensorimotor and social template, fostering a neuro-cultural signature that supports social attunement and identity formation within the DN. Segregation of the CON/DAN and the VAN into latent dimensions associated with community and family values, respectively, suggests differentiation of environmental (i.e., macro-ecological determinants) and affective (e.g., emotional or social cues) salience.

Critically, while these findings are restricted to the early adolescent window (ages 12–14 years), they underscore that the brain’s functional integration is an adaptive response to the unique sociocultural environment in which the child resides. For Hispanic youth, family ethnic identity and heritage culture may act as a vital neuroprotective buffer, stabilizing the internal self-system (DN) against the physiological and attentional taxes (CON/DAN) imposed by community-level structural disadvantage. Ultimately, these data suggest that promoting adolescent wellbeing requires more than individual-level intervention; it necessitates a biocultural approach that addresses structural inequities in community opportunity while fortifying the cultural integrity and linguistic traditions of the family unit.

## RESEARCH HIGHLIGHTS

- We examined multivariate associations between sociocultural environment at the individual, family, and community levels and resting state functional connectivity among Hispanic adolescents.
- Our results demonstrate a dual-pathway embedding, such that an ecological pathway links neighborhood opportunity and advantage to attentional systems and a cultural pathway links family ethnic identity and heritage culture retention to social-sensory systems.
- We established a multivariate link between macro-ecological stressors, large-scale brain connectivity and respiratory functioning, suggesting a systemic allostatic load.
- This work underscores the necessity of disparities-related cultural interventions that address structural inequities in child opportunity while fortifying the cultural integrity of the family unit.

## ACKNOWLEDGMENTS

Funding for this project was provided by NIH U01-DA041156 (ARL, MS, RG), R01-ES032295 (MMH), R01-ES031074 (MMH), K01-MD018069 (ET), and K01-DK135847 (SA). The Southern California Center for Latino Health (funded by the National Institute on Minority Health and Health Disparities, P50-MD017344) and The Saban Research Institute at Children’s Hospital of Los Angeles also provided support. Additional thanks to the FIU Instructional & Research Computing Center (IRCC, http://ircc.fiu.edu) for providing the HPC and computing resources that contributed to the research results reported within this paper.

Data used in the preparation of this article were obtained from the Adolescent Brain Cognitive DevelopmentSM (ABCD) Study (https://abcdstudy.org), held in the NIMH Data Archive (NDA). This is a multisite, longitudinal study designed to recruit more than 10,000 children age 9-10 and follow them over 10 years into early adulthood. The ABCD Study® is supported by the National Institutes of Health and additional federal partners under award numbers U01DA041048, U01DA050989, U01DA051016, U01DA041022, U01DA051018, U01DA051037, U01DA050987, U01DA041174, U01DA041106, U01DA041117, U01DA041028, U01DA041134, U01DA050988, U01DA051039, U01DA041156, U01DA041025, U01DA041120, U01DA051038, U01DA041148, U01DA041093, U01DA041089, U24DA041123, U24DA041147. A full list of supporters is available at https://abcdstudy.org/federal-partners.html. A listing of participating sites and a complete listing of the study investigators can be found at https://abcdstudy.org/consortium_members/. ABCD consortium investigators designed and implemented the study and/or provided data but did not necessarily participate in the analysis or writing of this report. This manuscript reflects the views of the authors and may not reflect the opinions or views of the NIH or ABCD consortium investigators. Additional support for this work was made possible from NIEHS R01-ES032295 and R01-ES031074.

## Competing Interests

The authors declare no competing interests.

## Author Contributions

ARL, KLB, and AM conceived and designed the project. CLH analyzed data. CLH, KS, KLB contributed scripts and pipelines. MMH and CC-I contributed methodology and resources. ARL and CLH wrote the paper and all authors contributed to the revisions and approved the final version.

## Notes

### Competing Interest Statement

The authors have declared no competing interest.

