## Supplemental Information for "Dual ecological-cultural pathways of sociocultural environment and resting state functional connectivity among Hispanic adolescents in the ABCD Study"

### SUPPLEMENTAL RESULTS

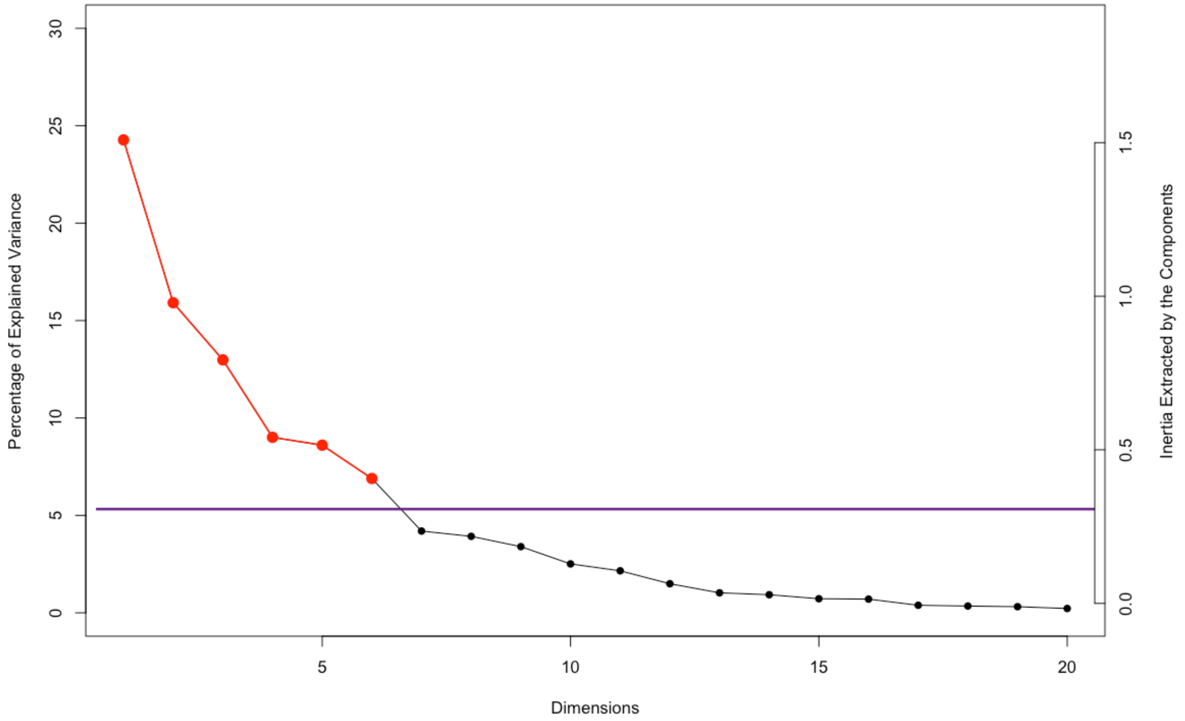


**Fig S1. PLSC Scree Plot.** Latent dimensions (y-axis) are plotted against the percentage of variance explained (x-axis). The horizontal reference line marks Kaiser's criterion; only the first three latent dimensions had both significant sociocultural and rsFC loadings. An elbow is observed at four latent dimensions, further justifying the retention of the first three latent dimensions, which cumulatively account for 53.17% of the shared variance.

| | Estimate (SE) | P Value | 95% CI Low | 95% CI High |
|---|---|---|---|---|
| **Latent Dimension 1** | | | | |
| **Physical Health** | | | | |
| BMI | -0.017 (0.011) | 0.132 | -0.039 | 0.005 |
| Blood Pressure | | | | |
| *Mean Systolic Pressure* | 0.005 (0.012) | 0.689 | -0.019 | 0.028 |
| *Mean Diastolic Pressure* | -0.009 (0.012) | 0.433 | -0.033 | 0.014 |
| Physical Activity | 0.008 (0.012) | 0.517 | -0.016 | 0.031 |
| Respiratory Functioning | | | | |
| *Wheezing/Whistling* | -0.039 (0.019) | 0.037* | -0.076 | 0.002 |
| *Dry Cough* | 0.004 (0.018) | 0.828 | -0.032 | 0.040 |
| *Asthma* | -0.008 (0.018) | 0.660 | -0.042 | 0.027 |
| *Pneumonia/Bronchitis* | -0.038 (0.033) | 0.251 | -0.104 | 0.027 |
| Sleep | | | | |
| *Average Weekly Duration* | 0.021 (0.012) | 0.082 | -0.003 | 0.045 |
| *Average Weekly Chronotype* | -0.004 (0.012) | 0.763 | -0.027 | 0.020 |
| **Mental Health** | | | | |
| Internalizing Behaviors | -0.014 (0.011) | 0.227 | -0.036 | 0.009 |
| Externalizing Behaviors | -0.001 (0.011) | 0.908 | -0.021 | 0.908 |
| **Latent Dimension 3** | | | | |
| **Physical Health** | | | | |
| BMI | 0.003 (0.008) | 0.694 | -0.020 | 0.694 |
| Blood Pressure | | | | |
| *Mean Systolic Pressure* | 0.006 (0.009) | 0.500 | -0.012 | 0.024 |
| *Mean Diastolic Pressure* | 0.012 (0.009) | 0.161 | -0.005 | 0.030 |
| Physical Activity | -0.006 (0.009) | 0.491 | -0.024 | 0.011 |
| Respiratory Functioning | | | | |
| *Wheezing/Whistling* | -0.010 (0.014) | 0.499 | -0.037 | 0.018 |
| *Dry Cough* | -0.005 (0.014) | 0.708 | -0.032 | 0.022 |
| *Asthma* | -0.011 (0.013) | 0.406 | -0.037 | 0.015 |
| *Pneumonia/Bronchitis* | -0.006 (0.025) | 0.800 | -0.055 | 0.043 |
| Sleep | | | | |
| *Average Weekly Duration* | -0.005 (0.009) | 0.550 | -0.023 | 0.012 |
| *Average Weekly Chronotype* | -0.012 (0.009) | 0.157 | -0.030 | 0.005 |
| **Mental Health** | | | | |
| Internalizing Behaviors | 0.011 (0.008) | 0.208 | -0.006 | 0.027 |
| Externalizing Behaviors | 0.003 (0.009) | 0.686 | -0.013 | 0.020 |
| * P < 0.05; ** P < 0.01 | | | | |

**Table S1. Latent Dimension Associations with Adolescent Physical and Mental Health.**
